# disperseNN2: a neural network for estimating dispersal distance from georeferenced polymorphism data

**DOI:** 10.1101/2023.07.30.551115

**Authors:** Chris C. R. Smith, Andrew D. Kern

## Abstract

Spatial genetic variation is shaped in part by an organism’s dispersal ability. We present a deep learning tool, `disperseNN2`, for estimating the mean per-generation dispersal distance from georeferenced polymorphism data. Our neural network performs feature extraction on pairs of genotypes, and uses the geographic information that comes with each sample. These attributes led `disperseNN2` to outperform a state-of-the-art deep learning method that does not use explicit spatial information: the mean relative absolute error was reduced by 33% and 48% using sample sizes of 10 and 100 individuals, respectively. `disperseNN2` is particularly useful for non-model organisms or systems with sparse genomic resources, as it uses unphased, single nucleotide polymorphisms as its input. The software is open source and available from https://github.com/kr-colab/disperseNN2, with documentation located at https://dispersenn2.readthedocs.io/en/latest/.

## 1 Background

The per-generation dispersal distance of an organism is a critical variable for the management of endangered and invasive species, understanding range shifts under climate change, and studying vectors of human disease [1, 2, 3]. A potent source of information that may be used to estimate this ecologically-relevant parameter is population genetic data that are geographically distributed. Accordingly, numerous methods to perform dispersal estimation have been proffered in the literature. For example, Rousset [4] presented a formula that estimates neighborhood size from the slope of the least squares fit of genetic distance against geographic distance. Dispersal rate can in turn be calculated from neighborhood size if the population density is also known. Rousset’s approach is currently the most widely used genetic-based method because it can be used with polymorphism data like short sequence repeats or single nucleotide polymorphisms (SNPs). Other dispersal estimation methods require very high-depth sequencing combined with statistical inference to obtain the necessary input data types. In particular, Ringbauer et al. [5] and Osmond and Coop [6] estimate dispersal rate using identity-by-descent blocks and genome wide inferred genealogies, respectively, and demonstrate their methods on taxa with exceptional genomic resources: humans and Arabidopsis. While approaches for inferring the latter data types are continually improving, they are still unavailable for most species.

We previously presented a deep learning tool, called `disperseNN`, that estimates dispersal rate using input data that are accessible even for some non-model species, SNPs, and that performs as well or better than existing methods [7]. Notably our previous method relied only on population genetic variation and the width of the sampling area; it did not utilize the spatial coordinates of individuals. In the current study, we present an improved neural network architecture, called `disperseNN2`, that explicitly uses geographic information and provides substantial performance gains over `disperseNN`, which was already more accurate than previous methods for small sample sizes.

## 2 Implementation

### 2.1 Overview

The `disperseNN2` program uses a deep neural network trained on simulated data to infer the mean, per-generation parent-offspring distance. Specifically, we aim to infer *σ*, the root-mean-square displacement along a given axis between a randomly chosen child and one of their parents chosen at random [4, 5]. `disperseNN2` is designed for SNP data obtained from reduced representation or whole genome sequencing, with either short-range or full linkage information. Because the model is trained on simulated data, the general workflow requires generating training datasets that accurately reflect the empirical genotypes of interest. While the neural network model has diverged substantially and is described below in detail, the general approach and analysis workflow are similar to Smith et al. [7]. The `disperseNN2` documentation includes complete instructions, example commands for the analysis workflow, and a number of usage vignettes (https://dispersenn2.readthedocs.io/en/latest/).

### 2.2 Network architecture

`disperseNN2` uses a pairwise convolutional network that performs feature extraction on *pairs* of individuals at a time (Figure 1). The first part of the model, which we refer to as “the extractor”, extracts pertinent information from pairs of genotypes, and merges the extracted features from all combinatorial pairs into a summary table for downstream processing. The latter part of the model uses the extracted data from many sample-pairs to predict *σ*. This strategy allows us to convey spatial information to the network, which is accomplished by attaching the geographic distance between each sample-pair directly to the genotype summaries from the corresponding pair.

**Figure 1.**
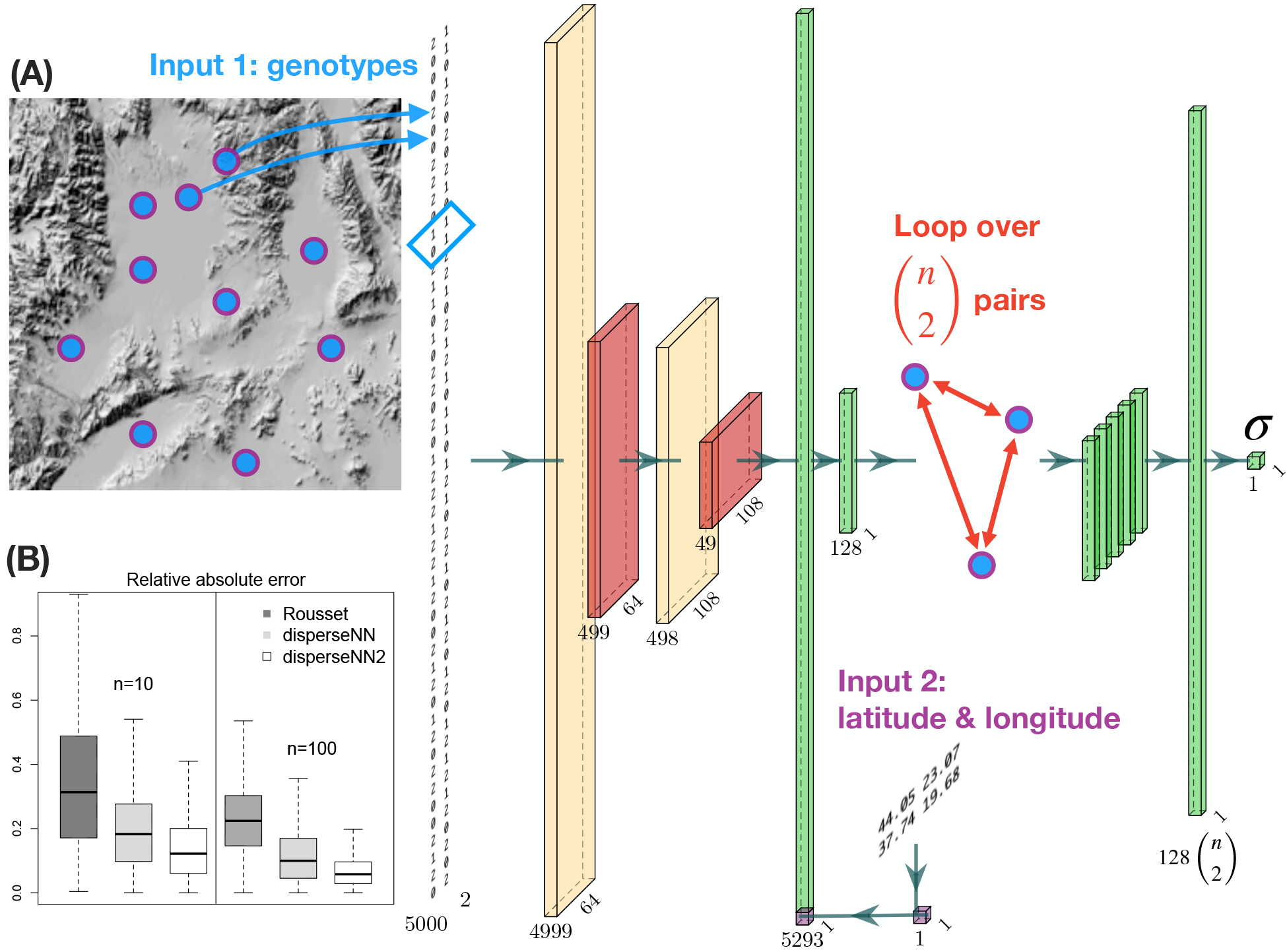
(A. neural network schematic) From left to right: a pair of individuals is selected for the feature-extraction step—this will be repeated for *k*_extract_ pairs. The genotype matrix shows the genotypes for the pair. Cream colored tensors are the output from convolution layers. The blue box over the genotypes shows the convolution kernel for the first layer. Red tensors are the output from pooling layers. The spatial coordinates for the current pair are subsetted from the locations table (Input 2). The Euclidean distance is concatenated with the flattened convolution output. Green tensors are the output from flattening, concatenating, or dense layers. The extractor is repeated for 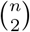 different pairs of individuals, and the resulting features are concatenated together. The dimensions noted beneath each tensor will vary depending on the input size; this example uses 5,000 SNPs (although the image of the genotypes shows a smaller number of SNPs). The visualized size of each tensor is proportional to the square root of the actual dimensions. Neural network images were generated using `PlotNeuralNet` (https://github.com/HarisIqbal88/PlotNeuralNet). (B. box plots) Also shown are validation results using Rousset’s method (dark grey), `disperseNN` (light grey), and `disperseNN2` (white), with two different sample sizes. Outliers are excluded.

The first input to `disperseNN2` is a genotype matrix consisting of minor allele counts (0s, 1s, and 2s) for *m* SNPs, ordered by genomic position, from *n* individuals. The program has the option to use unphased or phased genotypes; if phased, there are two genotypes (0s or 1s) per individual. However, rather than show the full genotype matrix to the network, we loop through all c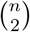 pairs of individuals and sub-set the genotypes of each pair. Feature extraction is then performed independently on each pair using convolution and pooling, where the convolution kernel spans two SNPs and each pooling step averages ten SNPs.

The second input is a table of geographic coordinates for the sample locations. As with the genotypes, the *x* and *y* coordinates are sub-set for each sample-pair. The Euclidean distance is calculated between the individuals within each pair and concatenated with the convolved genotype information for the pair. Last, the concatenated features are put through a fully connected layer, resulting in a vector of information gleaned from the pair. Weights are shared across all sample-pairs in the extractor.

After performing feature extraction on each pair of individuals, the features from all pairs are stacked together, and a final, fully connected layer with a single filter and linear activation is used to produce an estimate for *σ*. All other layers with trainable weights include rectified linear unit activation functions.

By default the network uses all combinatorial pairs of samples. However, GPU memory might become limiting with larger sample sizes. For example, with *n* = 100 there are 4950 sample-pairs. While one solution might be to omit some pairs from the analysis entirely, we instead exclude a number of pairs from the optimization of certain model parameters. Specifically, `disperseNN2` has the option to stop some pairs from contributing to the calculation of the gradient with respect to the weights in the extractor. Under this strategy, a smaller number of randomly chosen pairs, *k*_extract_, are used to optimize the extractor’s weights, which reduces memory demands appreciably. Meanwhile, features are still extracted from the full set of pairs during the forward pass, and all pairs help optimize the weights in the latter half of the network. In our validation analysis (see below) with *n* = 100, memory usage decreased linearly from 95.7 Gb with *k*_extract_ = 4950 to 12.0 Gb with *k*_extract_ = 100 (Gb= 0.017*k*_extract_+11.0). The ideal value for *k*_extract_ will likely depend on the total number of pairs, the number of SNPs, the batch size, and available memory. Tensorflow [8] and Keras (https://github.com/keras-team/keras) libraries were used to develop `disperseNN2`.

### 2.3 Generating training data

To use `disperseNN2`, researchers must simulate training datasets that are tailored to the study system. In particular, producing training data requires deciding on training distributions for dispersal rate and other parameters. Some other values that are relevant to dispersal inference, i.e. nuisance parameters, include population density, demographic history, and the shape and size of the species distribution [7]. Choices for these simulation settings will depend on the information available in each unique species or population, and if misspecified will increase error in dispersal predictions. However, the model may be trained to ignore certain nuisance parameters by varying them among the training simulations [7]. Recent developments, namely the programs `SLiM` [9] and `slendR` [10], make it feasible to simulate genomes in continuous space. The `disperseNN2` repository includes a `SLiM` script that may serve as a template for new simulations, and the `SLiM` manual includes general information for designing spatial simulations.

### 2.4 Analysis

The `disperseNN2` software includes tools for pre-processing training datasets, making it easy for practitioners to turn tree sequences (e.g., `SLiM` output) into `disperseNN2` input. Specifically, a sample of individuals is taken from the final generation of each simulation, and their genotypes and locations—inputs to the neural network—are saved in binary format to improve training efficiency. To mirror the empirical sampling scheme, individuals are chosen from simulations that are closest to the empirical sample localities, after projecting the empirical latitude and longitude onto a flat surface—this step aims to make the training data as similar to the empirical data as possible. The empirical data input to `disperseNN2` are unphased or phased SNP data in standard variant call format (VCF) and a corresponding table of latitude and longitude coordinates for each sample. The model is trained with mean squared error loss, Adam optimizer, and learning rate 10^*−*4^. The training duration depends on the input size: for example, using a training set of 50,000 datasets, each with 5,000 SNPs and *n* = 10, it takes 4.5 hours for 100 training iterations on a GPU; with *n* = 100 it takes approximately a week using all 4950 pairs and *k*_extract_ = 100. The program also parallelizes well across CPUs: using 50 cores leads to similar performance to one GPU. The `disperseNN2` documentation provides a complete vignette taking a user through the cycle of simulation, training, and eventual prediction for an empirical dataset.

## 3 Results

For benchmarking the new software we used simulated data as described in Smith et al. [7], using SLiM [9]. Briefly, the simulated genome is a single chromosome with length 10^8^ base pairs, with recombination rate 10^*−*8^ crossovers per base pair per generation. The habitat is a 50 *×* 50 square; local carrying capacity was set to 5 individuals per square map unit; and the mother-offspring dispersal distance, mating distance, and competition distances all shared the same value, *σ*_*f*_, which varied uniformly between 0.2 and 3. To obtain the “effective” dispersal rate to a randomly chosen parent, *σ*, the simulation parameter *σ*_*f*_ was multiplied by 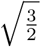 (see Smith et al., 2023). Importantly, this is a relatively simple model, without demographic perturbations or heterogeneous environment, designed only for benchmarking the new neural network. One hundred individuals were sampled uniformly at random at the end of the simulation. The full training set consisted of 1,000 SLiM simulations, each sampled 50 times, for a total of 50,000 training datasets.

We validated the `disperseNN2` architecture on 1000 held-out simulations and measured performance as the mean relative absolute error (MRAE), root mean squared error (RMSE), and the correlation between true and predicted values (*r*^2^). Prediction error with `disperseNN2` was substantially decreased relative to `disperseNN:` using 100 spatially distributed individuals and 5,000 SNPs as input, we observed a 48% reduction in MRAE with `disperseNN2` relative to `disperseNN` (Table 1). Using a smaller sample size, *n* = 10, we observed a 33% reduction in MRAE. This is a tremendous improvement to what was already state-of-the-art software. For reference, the MRAE with `disperseNN2` was 75% lower than Rousset’s method run on the same test data using *n* = 100, and 70% lower with *n* = 10. Furthermore, 1.9% and 22% of tests with Rousset’s method using *n* = 100 and *n* = 10, respectively, produced undefined output and were not included in the error calculation. See Smith et al. [7] for details on our implementation of Rousset’s method. In the *n* = 100 experiment we used all 4950 pairs, but found that using *k*_extract_ = 100 instead of *k*_extract_ = 4950 reduced memory consumption and computation time considerably, without a reduction in accuracy.

**Table 1:** Error metrics.

| Method | Sample size | MRAE | RMSE | $r^2$ |
| --- | --- | --- | --- | --- |
| **disperseNN2** | 10 | 0.140 | 0.440 | 0.833 |
| **disperseNN** | 10 | 0.208 | 0.558 | 0.708 |
| Rousset | 10 | 0.458 | 3.318 | 0.058 |
| **disperseNN2** | 100 | 0.065 | 0.152 | 0.980 |
| **disperseNN** | 100 | 0.124 | 0.368 | 0.852 |
| Rousset | 100 | 0.255 | 1.086 | 0.425 |

Importantly, these tests used data that were generated under the same process that produced the training data, and only a single model parameter, *σ*, was unknown. Thus, the reported accuracy represents a best case scenario. In practice, there may be additional unknown parameters, for example, population density, that should be incorporated into training by varying the unknown parameters between simulations (i.e., using a ‘prior’ distribution). See Smith et al. [7] for further discussion and experiments involving model misspecification.

## 4 Conclusion

We present a novel deep learning architecture, `disperseNN2`, for estimating the mean, per-generation dispersal rate from genotypes and their geographic coordinates. The `disperseNN2` neural network differs from our previous model, `disperseNN`, in two ways. First, `disperseNN2` loops through *pairs* of genotypes at a time, extracting relevant information from each pair. Second, the neural network makes use of the geographic coordinates associated with each genotype. These changes allow `disperseNN2` to outperform `disperseNN` by a sizeable margin. Our approach will be especially useful for non-model organisms that lack accurate identity-by-descent tracts and genome wide genealogies, because it can be used with unphased SNP data. One limitation for our method is generating the required training data, which must be designed carefully to reflect the empirical data of interest and can be computationally expensive for large populations.

The effective *σ* parameter output by `disperseNN2` represents a measure of gene flow across space over generations. Inferring this critical evolutionary parameter for a species allows modeling of a number of affected phenomena, for example, the spread of an adaptive allele in a population [11], or, for measuring the strength of selection against hybrids in a genomic cline [12]. Further, we may learn about the evolution of dispersal by comparing *σ* between taxa, or by regressing *σ* with environmental variables. Effective *σ* has contributions from both the mother-offspring distance (e.g., seed dispersal) and the mating distance (e.g., pollination distance). Therefore, if either the effective mother-offspring or effective mating distance is known for a species, the other can be inferred using genetics-based methods like ours. In other cases, it might be reasonable to assume the two distances occur on similar scales. This ecological information can in turn be used to study habitat connectivity, guide conservation translocations, and to predict species range shifts.

Whereas some studies in population genetics have applied convolutional neural networks to the full genotype matrix, our deep learning model performs feature extraction on pairs of genotypes. Having the network focus on pairs is an intuitive strategy for many genetics applications, particularly those involving spatial genetic data, where researchers are often interested in the relatedness between individuals. For studying dispersal, this approach brings the model’s attention to the genetic and geographic distances between individuals, which follows the basic strategy of well-established models like that of Rousset [4]. Architectures like this one may be useful for explicitly between-individual tasks like characterizing identity-by-descent tracts, or for other tasks in population genetics like detecting selective sweeps or inferring demographic history.

## 5 Abbreviations

*MRAE*: mean relative absolute error
*RMSE*: root mean squared error
*SNP*: single nucleotide polymorphism
*VCF*: variant call format

## 6 Declaration

### Ethics approval and consent to participate

Not applicable.

#### Consent for publication

Not applicable.

#### Availability of data and materials

*Project name*: disperseNN2.

*Project home page*: https://github.com/kr-colab/disperseNN2.

*Operating system(s)*: Platform independent.

*Programming language*: Python.

*Other requirements*: None.

*License*: MIT.

*Any restrictions to use by non-academics* : None.

#### Funding

This work was supported by the National Institutes of Health awards F32GM146484 to C.S. and R01HG010774 and R35GM148253 to A.D.K..

## Acknowledgements

We thank Peter Ralph and members of the Kern-Ralph colab for comments on the project and manuscript.

## Authors’ contributions

C.S. and A.K. conducted research and wrote the manuscript.

## References

[1] Don A Driscoll, Sam C Banks, Philip S Barton, Karen Ikin, Pia Lentini, David B Lindenmayer, Annabel L Smith, Laurence E Berry, Emma L Burns, Amanda Edworthy, et al. The trajectory of dispersal research in conservation biology. Systematic review. PloS one, 9(4):e95053, 2014.

[2] Catriona M Harris, Kirsty J Park, Rachel Atkinson, Colin Edwards, and Justin MJ Travis. Invasive species control: incorporating demographic data and seed dispersal into a management model for Rhododendron ponticum. Ecological Informatics, 4(4):226–233, 2009.

[3] James Orsborne, Luis Furuya-Kanamori, Claire L Jeffries, Mojca Kristan, Abdul Rahim Mohammed, Yaw A Afrane, Kathleen O’Reilly, Eduardo Massad, Chris Drakeley, Thomas Walker, et al. Investigating the blood-host plasticity and dispersal of Anopheles coluzzii using a novel field-based methodology. Parasites & vectors, 12(1):1–8, 2019.

[4] Francois Rousset. Genetic differentiation and estimation of gene flow from F-statistics under isolation by distance. Genetics, 145(4):1219–1228, 1997.

[5] Harald Ringbauer, Graham Coop, and Nicholas H Barton. Inferring recent demography from isolation by distance of long shared sequence blocks. Genetics, 205(3):1335–1351, 2017.

[6] Matthew M Osmond and Graham Coop. Estimating dispersal rates and locating genetic ancestors with genome-wide genealogies. bioRxiv, pages 2021–07, 2021.

[7] Chris CR Smith, Silas Tittes, Peter L Ralph, and Andrew D Kern. Dispersal inference from population genetic variation using a convolutional neural network. Genetics, 224(2):iyad068, 2023.

[8] MartÍn Abadi, Ashish Agarwal, Paul Barham, Eugene Brevdo, Zhifeng Chen, Craig Citro, Greg S Corrado, Andy Davis, Jeffrey Dean, Matthieu Devin, et al. Tensorflow: Large-scale machine learning on heterogeneous distributed systems. arXiv preprint arXiv:1603.04467, 2016.

[9] Benjamin C Haller and Philipp W Messer. SLiM 3: forward genetic simulations beyond the Wright-Fisher model. Molecular biology and evolution, 36(3):632–637, 2019.

[10] Martin Petr, Benjamin C Haller, Peter L Ralph, and Fernando Racimo. slendr: a framework for spatio-temporal population genomic simulations on geographic landscapes. bioRxiv, pages 2022–03, 2022.

[11] Margaret C Steiner and John Novembre. Population genetic models for the spatial spread of adaptive variants: A review in light of sars-cov-2 evolution. PLoS Genetics, 18(9):e1010391, 2022.

[12] Nicholas H Barton and Brian Charlesworth. Genetic revolutions, founder effects, and speciation. Annual review of ecology and systematics, 15(1):133–164, 1984.

